# Acute oral toxicity and risks of exposure to the neonicotinoid thiamethoxam, and other classes of systemic insecticide, for the Common Eastern Bumblebee (*Bombus impatiens*)

**DOI:** 10.1101/2020.01.27.921510

**Authors:** Kayla A. Mundy-Heisz, Ryan S. Prosser, Nigel E. Raine

## Abstract

The Common Eastern Bumblebee (*Bombus impatiens*) is native to North America with an expanding range across Eastern Canada and the USA. This species is commercially produced primarily for greenhouse crop pollination and is a common and abundant component of the wild bumblebee fauna in agricultural, suburban and urban landscapes. However, there is a dearth of pesticide toxicity information about North American bumblebees. The present study determines the acute oral toxicity (48-hour LD50) of cyantraniliprole (>0.54 μg/bee), flupyradifurone (>1.7 μg/bee), sulfoxaflor (0.0194 μg/bee), and thiamethoxam (0.0012 μg/bee). Compared with published honey bee (*Apis mellifera*) LD50 values, the present study shows that thiamethoxam and sulfoxaflor are 4.2x and 7.5x more acutely toxic to *B. impatiens*, whereas flupyradifurone is more acutely toxic to *A. mellifera*. The current rule of thumb for toxicity extrapolation beyond the honey bee as a model species, termed 10x safety factor, may be sufficient for bumblebee acute oral toxicity. A comparison of three risk assessment equations suggested that the Standard Risk Approach (SRA) and Fixed Dose Risk Approach (FDRA) provide more nuanced levels of risk evaluation compared to the European Plant Protection Organization (EPPO) Hazard Quotient (HQ), primarily because SRA and FDRA take into account real world variability in pollen and nectar pesticide residues and the chances that bees are exposed to them.

## 1 INTRODUCTION

The pollination services provided by wild bees are important for maintaining the integrity of wild plant communities and play a key role in the agricultural production of crops, including fruits, vegetables and nuts (Klein et al. 2007). However, wild pollinators are not considered in the risk assessment process, even though they are present in agricultural landscapes. Currently, the European honey bee (*Apis mellifera*) is the only required model species for regulatory ecotoxicological risk assessments for all insect pollinators (European and Mediterranean Plant Protection Organization 2010). The risk assessment scaffold starts with acute oral and contact toxicity testing to establish lethal endpoints (such as the exposure level that kills 50% of individuals within a given time period, i.e. the LD50) within a laboratory setting (i.e., tier 1 assessments). Demonstrating high levels of toxicity in these tests can trigger higher tier testing, culminating with (tier 3) field studies that are realistic to the proposed use(s) of the chemical under consideration and most frequently use assessments of honey bee colony strength as effect endpoints (European and Mediterranean Plant Protection Organization 2010). This scaffold assessment has deficiencies depending on the scale of the toxicity tests, for example, third tier field assessments using honey bee hives containing 5000-10000 individuals are unlikely to be representative of a bumblebee colony containing a few hundred individuals (Gradish et al. 2019) or solitary bees, which are solo provisioners (Sgolastra et al. 2019). Furthermore, the wide diversity in ecology and life history of bee species are not taken into consideration when warning labels or best management practices are developed for pesticide products (European and Mediterranean Plant Protection Organization 2010). For example, risk management statements might recommend that pesticide users “remove or cover beehives during application” (European and Mediterranean Plant Protection Organization 2010, pg. 326) – advice that is only relevant for managed honey bees or bumblebees, whilst overlooking wild bees.

Pesticide regulators have begun to consider expanding the scope of ecotoxicological risk assessments beyond the honey bee as the model species for insect pollinators. These developments have been driven by a rapidly expanding body of evidence of impacts of exposure to pesticides from non-*Apis* bee species (Godfray et al. 2014; Godfray et al. 2015; Lundin et al. 2015), and concerns that pesticide sensitivity can vary substantially among bee species (Arena and Sgolastra 2014; Gradish et al. 2019; Rundlöf et al. 2015; Sgolastra et al. 2019; Woodcock et al. 2017). Notably, many of these studies have examined the potential impacts of neonicotinoid insecticide exposure on bumblebees, particularly the European commercially-reared species *Bombus terrestris* (Rundlöf et al. 2015; Woodcock et al. 2017). This highlights the need to fully assess any variability in sensitivity among bee taxa to the different types of pesticides to which they could be exposed.

Neonicotinoids are currently the most widely used class of insecticide around the world (Jeschke et al. 2011). These systemic insecticides are frequently applied as seed treatments or soil applications in agricultural crops. Systemic insecticides are highly water-soluble allowing absorption from the soil into tissues of the crop plant (Simon-Delso et al. 2015). This can provide effective protection against pests that damage plants by piercing the stem and those that consume other plant parts (Sparks et al. 2013). However, high water solubility also leads to movement through the soil profile and into non-target water courses. These insecticides may be found in the pollen and nectar of the plant, and if the crop requires pollination for fruit set, non-target insects including pollinators, may come into contact with these residues as they visit flowers and consume these floral rewards (Bonmatin et al. 2015; Godfray et al. 2014; Chagnon et al. 2015). Systemic insecticides have also been detected in non-crop flowers (David et al. 2016; Long and Krupke 2016), increasing the potential for non-target insect exposure.

The mode of action of these insecticides plays a key role in their toxicity to both pest and non-target insects. Neonicotinoids affect insects by binding to the acetylcholine site of the nicotinic acetylcholine receptor (nAChR) modulating or stopping the passage of information through the central nervous system and producing symptoms ranging from hyper-excitation to paralysis and death (Simon-Delso et al. 2015). Since their introduction in the early 1990s there has been increasing evidence of unintended negative consequences of their use for bees (Godfray et al. 2014; Godfray et al. 2015; Lundin et al. 2015). Neonicotinoids may be becoming less effective at controlling target pests as tolerance and resistance have been documented for the brown planthopper (Liu et al. 2005), and evidence of economic benefits for farmers are less clear cut (Krupke et al. 2017; Mourtzinis et al. 2019). Concerns about impacts to non-target organisms have already led to use restrictions of neonicotinoids in some regions of the world, with more regulatory changes likely to follow. For example, clothianidin, imidacloprid and thiamethoxam are no longer available for use on crops grown outside greenhouses in the European Union (European Union 2013; 2018a,b,c) and reductions in usage of these insecticides have been mandated in Ontario, Canada since 2015 (Ontario Ministry of Agriculture Food and Rural Affairs 2015). While these regulatory changes have been occurring to restrict neonicotinoid usage, other classes of systemic insecticides are being introduced and used for pest control as alternatives or replacements for neonicotinoids.

The sulfoximine class, represented by the sole active ingredient sulfoxaflor, has already been identified as a potential threat to pollinators (Brown et al. 2016; Siviter et al. 2018) due to the shared mode of action with neonicotinoids, via the nAChRs. Butenolides are another insecticide class with this mode of action, but with a different pharmacophore system, represented by the sole active ingredient flupyradifurone (Insecticide Resistance Action Committee 2019; Jeschke et al. 2015). In contrast, diamide insecticides, such as chlorantraniliprole and cyantraniliprole (Insecticide Resistance Action Committee 2019), paralyse insects by affecting their nerves and muscles. Diamides act on the ryanodine receptors responsible for intracellular calcium store regulation within the sarcoplasmic reticulum of insect muscle cells (Lahm et al. 2005). Thus, while systemic insecticides vary in mode of action, they are all highly water solubility and can therefore move through soil and potentially be taken up by non-target plants.

Neonicotinoids, butenolides, sulfoximines and diamides are all classes of systemic insecticides to which non-target insects may become exposed, have been registered for use following a honey bee centric risk assessment scaffolding, and if there is research regarding impacts for insect pollinators it is skewed towards honey bees and *Bombus terrestris* in Europe. To address knowledge gaps for North American pollinators, the acute oral toxicity to Common Eastern Bumblebee (*Bombus impatiens*) workers was compared among representatives from these four systemic insecticide classes: thiamethoxam (neonicotinoid), flupyradifurone (butenolide), sulfoxaflor (sulfoximine) and cyantraniliprole (diamide). Results from these bumblebee tests were compared to published acute oral toxicity data of these insecticides for the European honey bee (*Apis mellifera*), and an assessment of the potential hazard of these insecticides to *B. impatiens* workers was conducted.

## 2 MATERIALS AND METHODS

### 2.1 Test organisms

Seven research colonies (*Bombus impatiens* Cresson), each containing 40-60 workers, a queen, and no males, were obtained from Biobest (Leamington, Ontario, Canada). Colonies were kept at 24 ± 1°C, 30% relative humidity and under constant darkness. Colonies were fed every other day with 3 grams of honey bee collected pollen (University of Guelph Honey bee Research Centre, Guelph, Ontario, Canada) and an artificial nectar substitute (Biobest, Leamington, Ontario, Canada) was provided *ad libitum*. Colonies were allowed to acclimatize for 24 hours before the workers were removed for trials.

### 2.2 Chemicals and analytical methods

Technical grade (98% pure) cyantraniliprole, flupyradifurone, sulfoxaflor and thiamethoxam were purchased from Toronto Research Chemicals (Toronto, Ontario, Canada). Previous work with range finding trials using five research colonies determined the nominal concentrations to be used in definitive testing. The highest concentrations for cyantraniliprole and flupyradifurone were determined based on maximum solubility in 50% sugar water at room temperature (23°C). Stock solutions were made by dissolving the active ingredient (A.I.) in 50% weight/volume sugar solution. We made initial stock solutions of 0.2 mg/mL for both cyantraniliprole and flupyradifurone, 0.04 mg/mL for sulfoxaflor, and 0.004 mg/mL for thiamethoxam. We created a series of different exposure solutions by performing serial dilutions of the initial stock solution for each insecticide. Dilutions for cyantraniliprole and flupyradifurone followed a 0.8 descending nominal geometric scale, whereas sulfoxaflor and thiamethoxam followed a 0.5 descending nominal geometric scale (Table 1).

**Table 1.** Nominal and measured concentrations of systemic insecticides from sugar solutions containing cyantraniliprole, flupyradifurone, sulfoxaflor or thiamethoxam.

| Active Ingredient | Nominal concentration (mg/mL) | Measured concentration (mg/mL) | Difference (%) |
| --- | --- | --- | --- |
| Cyantraniliprole | 0.2 | 0.054 | 73 |
|  | 0.16 | 0.049<sup>a</sup> | 69 |
|  | 0.128 | 0.044 | 66 |
|  | 0.102 | 0.008 | 92 |
|  | 0.082 | 0.005 | 94 |
| Flupyradifurone | 0.2 | 0.17 | 15 |
|  | 0.16 | 0.143<sup>a</sup> | 11 |
|  | 0.128 | 0.12 | 6 |
|  | 0.102 | 0.075 | 27 |
|  | 0.082 | 0.052 | 37 |
| Sulfoxaflor | 0.005 | 0.0089 | -78 |
|  | 0.0025 | 0.0049<sup>a</sup> | -95 |
|  | 0.00125 | 0.0027<sup>b</sup> | -112 |
|  | 0.000625 | 0.0014<sup>a</sup> | -129 |
|  | 0.0003125 | 0.0008 | -146 |
| Thiamethoxam | 0.001 | 0.00120 | -20 |
|  | 0.0005 | 0.00074 | -48 |
|  | 0.00025 | 0.00029<sup>a</sup> | -14 |
|  | 0.000125 | 0.00010 | 20 |
|  | 0.0000625 | 0.00006 | 7 |

<sup>a</sup> Calculated using the mean percentage difference of the nominal vs. measured concentration immediately before and the nominal vs. measured concentration immediately afterwards. For example, the cyantraniliprole nominal concentration of 0.16, has a “measured concentration” of 0.049, but this is an estimate based upon the average difference between the immediately before nominal concentration, 0.2 mg/mL, percent difference (73%) from the measured concentration, 0.054 mg/mL, and the immediately after nominal concentration, 0.128 mg/mL, percent difference (66%) from the measured concentration, 0.044 mg/mL.
<sup>b</sup> Calculated using the mean of highest nominal vs. measured concentration and lowest nominal vs. measured concentration percentage difference for that active ingredient.

Samples of each exposure solution were sent for analysis at the Agriculture and Food Laboratory (AFL, Guelph, Ontario, Canada). AFL is ISO/IEC 17025 certified and is recognized as a Good Laboratory Practice laboratory by the Standards Council of Canada. A multi-residue screen test was used to detect presence and quantity of active ingredient in each sample. Thus, ensuring only the intended insecticide was present and no cross contamination or mixtures occurred. Cyantraniliprole, flupyradifurone and thiamethoxam had four of the geometric dose samples submitted for confirmation. Sulfoxaflor had the highest and lowest concentration tested.

Briefly, 1 mL of each sample was diluted in 50 mL of water following the Quick, Easy, Cheap, Effective, Rugged, and Safe (QuEChERS) method. Aliquots of 20 mL were extracted into 1% acetic acid in acetonitrile in the presence of anhydrous sodium acetate and magnesium sulphate. The supernatant was evaporated and diluted with methanol and 0.1 M ammonium acetate. Sample extracts were analyzed using liquid chromatography coupled with electrospray ionization tandem mass spectrometry (LC/ESI-MS/MS). This methodology produced a limit of detection (LOD) of 2.0 x 10^-5^ mg of insecticide/mL of sugar solution and a limit of quantification (LOQ) of 6.0 x 10^-5^ mg/mL (Table 1).

### 2.3 Oral toxicity bioassay

To assess the toxicity of oral exposure to systemic insecticides workers were removed from colonies and quasi-randomly assigned to a treatment group (one of five insecticide concentrations or an untreated control), this was to ensure worker bees from all colonies were allocated to each treatment group to mitigate any potential colony level effects. A group of 8-12 bees were allocated to each insecticide concentration, and each test was repeated 4 times (Supplemental Table S1 and S2). Individual bees were transferred under red light into individual Titan 60-mL medicine cups. Once in the containers, bees were starved for 3-4 hours. Bees were assessed after the 3-hour starvation period by touching the cup to assess whether they were suitable for inclusion in the test. If individuals were dead or showed abnormal behaviour (including no defensive response when the cup was touched, a lack of buzzing or leg raising, slow movement, spinning in smaller circles on the spot, falling and remaining on back when attempting to climb the side of the cup) they were excluded from the bioassay. Bees included in the assay after starvation were then given 10 μL of either a treatment (sugar solution containing pesticide) or control (sugar only) solution, which was deposited on the bottom of the cup. Each bee had 20 minutes to completely consume the sugar solution to ensure all bees used in the bioassay received the same exposure within each treatment group. Any bees that failed to consume the entire 10 μL of solution were excluded from the bioassay. After consuming the sugar solution bees were moved into a growth chamber at 25 ± 1°C with a relative humidity between 50-60%. Each bee received 50% weight/volume (w/v) sugar solution via a soaked cotton bud (inserted through a single small X-shaped slit cut in the side of the cup), which was replenished every ~24 hours when they were checked for mortality. Those that did not produce a defence response via a leg lift were gently touched with forceps, if no movement or vibration was observed, bees were scored as dead. Mortality was assessed at 4, 24, 48, and 72 hours after treatment sugar solution consumption. Control mortality was < 20% across all trials (Table S2).

### 2.4 Sublethal effects

Sublethal effects that could be indicative of neurotoxic impacts were recorded at each assessment time. Sublethal effects included an inability to stand or walk upright, an inability to right itself from lying on its back, antennal and/or leg spasms, no defence response when the cup was touched, either a lack of buzzing or leg raising, slow movement, spinning in smaller circles on the spot, falling and remaining on back when attempting to climb the side of the cup. Upon initial inspection of movement to assess mortality those displaying sublethal effects were watched for 1 minute to confirm abnormalities in their behaviour, and those not moving were inspected for death. We calculated the proportion of bees demonstrating abnormal behaviours at each time point (4, 24, 48, 72 hours), excluding dead individuals, for each dose of each insecticide.

### 2.5 Honey bee (Apis mellifera) LD50 data

The Pest Management Regulatory Agency (PMRA) within Health Canada (Health Canada 2013; 2014; 2015) and US EPA’s ECOTOX (United States Environmental Protection Agency 2019) databases were used to find 48-hour oral LD50 data for honey bees (*Apis mellifera*) exposed to cyantraniliprole, flupyradifurone, sulfoxaflor and thiamethoxam.

### 2.6 Risk assessment

The traditional initial tier risk calculation published by the European Plant Protection Organization (EPPO) (European and Mediterranean Plant Protection Organization 2010) is the hazard quotient (HQ) (Equation 1). The HQ considers the amount of active ingredient (A.I.) within a commercial product (g A.I./ha) applied to the crop and the acute LD50 of that active ingredient (Equation 1). An HQ less than 50 represents a low-risk, between 50 and 2500 a moderate-risk, and more than 2500 a high-risk (European and Mediterranean Plant Protection Organization 2010). A value less than 50 at this stage would lead to the classification of low-risk to bees and the risk assessment would not proceed beyond the first tier (European and Mediterranean Plant Protection Organization 2010). If the HQ was greater than or equal to 50, a tier two, semi-field trial would subsequently be required (European and Mediterranean Plant Protection Organization 2010).

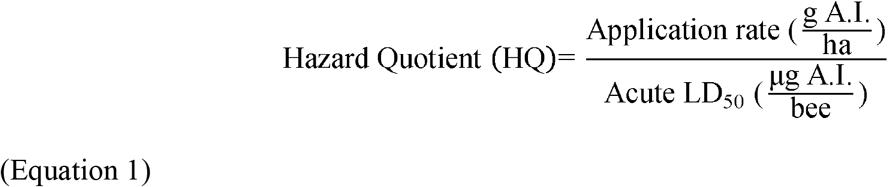

The minimum and maximum application rates from the manufacturer labels were determined for five commercial formulations of cyantraniliprole (Benevia^®^, Exirel^™^, Okina Insect Control, Verimark^®^, HGW86 200 SC Potato Seed Treatment Insecticide - all Dupont^™^ products), one for flupyradifurone (Sivanto^™^ Prime Insecticide), two for sulfoxaflor (Twinguard^™^ and Closer^™^ Insecticide), and one for thiamethoxam (Minecto^™^ Duo 40WG: Table S3). The mean, minimum, and maximum application rates were then input into the HQ equation (Equation 1) with the corresponding LD50 values determined in this study for each A.I. (Table S3). The outputs were then classified using EPPO guidelines as low, moderate, or high-risk to *B. impatiens* workers.

The proportion of low-, moderate- or high-risk scenarios was compared because a different number of HQ calculations were performed for each A.I. (Table S3). For each active ingredient, the total count of low (HQ <50), moderate (HQ = 50-2500) or high-risk (HQ >2500) hazard quotients was divided by the total number of HQs. For example, 163 HQs were calculated for the mean application rate of cyantraniliprole of which 161 (99%) were categorized as a moderate-risk and 2 (1%) were classified high-risk.

The standard risk approach (SRA) and the fixed dose risk approach (FDRA) were also applied using the data generated from this study. The SRA (Sanchez-Bayo and Goka 2014) includes the frequency with which a bee comes into contact with a pesticide residue (Equation 2). Thus, the SRA evaluates the probability of a pesticide residue causing 50% mortality among bees that come into contact with contaminated pollen or nectar resources (Sanchez-Bayo and Goka 2014). The value used in the risk calculation was the median frequency of detection for 160 pesticides, this ensured a standardized value across each pesticide tested (Sanchez-Bayo and Goka 2014). Thiamethoxam was the only insecticide in the present study to have a distinct frequency value reported (Sanchez-Bayo and Goka 2014), so these values were used in the SRA equations for thiamethoxam and the median frequencies were used for cyantraniliprole, flupyradifurone and sulfoxaflor. The likelihood of exposure values exceeding 5% are considered high-risk, 1-5% moderate-risk, and below 1% low-risk (Sanchez-Bayo and Goka 2014).

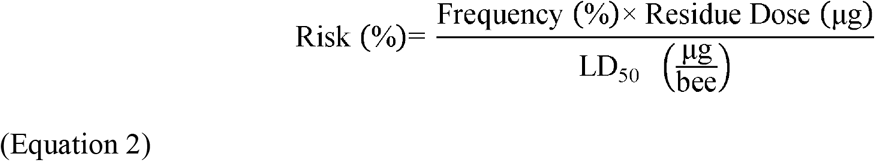

Sanchez-Bayo and Goka (2014) created the fixed dose risk approach (FDRA) to better encapsulate risk posed from chronic, dietary exposure (Equation 3). The FDRA estimates the time it takes for a bee to reach the LD50 based upon the estimated daily dose. Fewer than 2 days is considered high-risk, between 2 and 7 days a moderate-risk, and 7 or more days is considered low-risk (Sanchez-Bayo and Goka 2014).

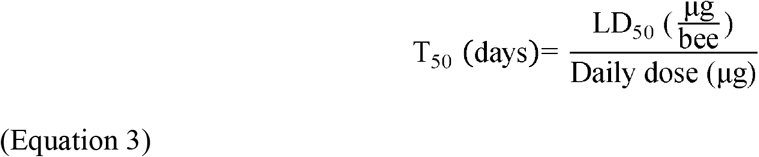

The SRA and the FDRA equations both used the same residue data to estimate exposure (compiled from Botías et al. 2015; David et al. 2016; Health Canada 2013; 2014; 2015; Tsvetkov et al. 2017), as commercial formulation application rates are not suitable for these equations (Table S4). As with the HQ, each active ingredient had a different number of calculations performed, due to the different number of approved products and their respective crop uses, so the proportion of low-, moderate- or high-risk values were compared.

### 2.7 Statistical analyses

RStudio Version 1.1.383 with R Version 3.3.2 (R Core Team 2019) with packages; dose response analysis (drc) (Ritz et al. 2015) and graphics (ggplot2) (Wickham 2009) were used to conduct statistical analysis and present data.

To characterize the dose-response relationship for each insecticide, a hierarchical model structure of the general four-parameter log-logistic function was used (Equation 4). As the data were binomial (alive = 0, dead = 1), the upper and lower limits were fixed as 0 and 1 while the model estimated the exposure that would lead to 50% mortality (LD50) (parameter e) and the relative slope (parameter b). This model was used to estimate an LD50 for each insecticide after 24, 48, and 72 hours post consumption based upon measured concentrations.

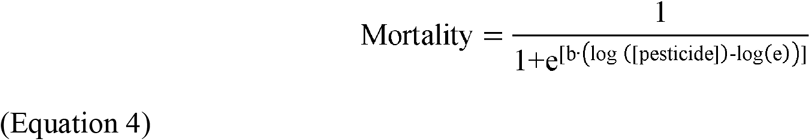

All active ingredient dose-response curves were compared using two models: model 1 assumed different LD50 values for each active ingredient, and model 2 assumed a common LD50 amongst all active ingredients. The common LD50 model (model 2) was created by adding in the *pmodels* argument to the original different LD50 model (model 1) (Ritz et al. 2015). The creation of different and a common LD50 for the active ingredients was repeated at three time points (24, 48, and 72 hours). The common and different LD50 models were compared using a likelihood ratio test, which is completed by the ANOVA function listing the common (all active ingredients have the same LD50) then the different LD50 model (active ingredients have different LD50s). If the likelihood ratio test value is significant, then the common and different LD50 models are not the same, indicating different LD50 values for each active ingredient and that confidence intervals for the LD50 for each active ingredient can be obtained.

### 2.7.1 Sublethal effects

A Shapiro-Wilk normality test was used to determine whether either measured exposure or the proportion of bumblebees displaying a sublethal response followed a normal distribution. We examined the association between the measured exposure and the proportion of bumblebees displaying a sublethal response using a Kendall Rank correlation coefficient.

### 2.7.2 Mean mass comparisons

Overall differences in the mean mass (g) of bumblebee workers belonging to a particular colony or assigned to a given treatment group were tested using either a Kruskal-Wallis test or ANOVA. A Wilcoxon test or Student’s t-test with a Bonferroni correction were used to compare the mean mass (g) of bumblebee workers among colonies and treatment groups.

## 3 RESULTS AND DISCUSSION

### 3.1 Test organisms

A total of 962 bumblebee workers were exposed to the test solutions of which 896 consumed the solution prior to the 20-minute cut off time. The percentage of bees that consumed test solutions was relatively high across active ingredients, trials, and doses (Table S1). For the four definitive tests, each insecticide had a minimum of 213 (range 213-242) bumblebees tested across five doses and a control. Mortality at the 48-hr checkpoint ranged from 3-13% for the control group and from 41-100% at the highest dose across insecticides (Table S2).

### 3.2 Chemical analysis

Analyses revealed differences between nominal and measured levels of insecticides found in serial dilutions used for exposure solutions in oral toxicity assays. On average measured concentrations were either 79% or 19% less than nominal concentrations for cyantraniliprole and flupyradifurone respectively (Table 1). Measured concentrations were on average 112% or 11% higher than nominal concentrations for sulfoxaflor and thiamethoxam respectively.

### 3.3 Oral toxicity bioassay

The random removal of bumblebees from colonies is a good way to ensure that intra- and inter-colony variation in worker mass is taken into consideration. In this study, the maximum difference in the mean mass of bumblebees tested within each active ingredient was 28 mg. Comparing the mean mass of workers used in the trials from the seven test bumblebee colonies demonstrated an overall significant difference (*p* = 0.026, Figure S1). Colony K (0.138 ± 0.038 g) contained significantly larger workers than both colonies H (0.119 ± 0.041g, *p* <0.01) and J (0.121 ± 0.040, *p* < 0.05) when directly compared using Wilcoxon tests with Bonferroni correction (Figure S1).

No significant differences in mean mass of workers that consumed or did not consume test solutions within twenty minutes (*p* = 0.51) were found, but the mean mass of workers differed significantly across the four active ingredients tested (*p* < 0.001, Kruskal-Wallis; Figure S2). Comparing the mean mass of bumblebees among the four repeated trials for each active ingredient, only significant differences for cyantraniliprole were detected (*p* < 0.001, Wilcoxon; Table S5, S6). The mean mass of workers allocated for each treatment dosage were significantly different for sulfoxaflor (*p* = 0.016, Kruskal-Wallis; Table S5, S7), but there was only a significant difference between the sulfoxaflor doses 0.0143 and 0.089 μg A.I./bee (*p* = 0.04, Wilcoxon; Table S7).

All active ingredients had a different LD50 values at 24, 48 and 72 hours (Table 2). The resultant LD10 and LD25 were also different at the 48-hr time point (Table S8). The assumption of different LD50 values for the four active ingredients was the most accurate model (*p* < 0.001) and used to estimate the LD50 (parameter e) and slope (parameter b) for each active ingredient. The dose response curve at 48 hours for flupyradifurone was significantly different from sulfoxaflor (*p* < 0.001), but not thiamethoxam (*p* = 0.123). Curves for sulfoxaflor and thiamethoxam were also significantly different (*p* < 0.001). The estimated relative potency based upon the LD50 at 48 hours shows thiamethoxam to be the most potent, followed by sulfoxaflor and flupyradifurone. Sulfoxaflor was approximately half as potent as thiamethoxam.

**Table 2.** LD_50_ (parameter e, μg/bee) and slope (parameter b) estimates of four active ingredients at three time points (24, 48, and 72 h) for *Bombus impatiens* workers. CYA failed to fit to a dose response model at 48 hours, and the sample size was insufficient for 72 hours.

| 24 hour |  |  |  |  |  |  |  |
| --- | --- | --- | --- | --- | --- | --- | --- |
|  | LD50 |  |  |  | Slope |  |  |
|  | EST | S.E. | LOW | UPP | EST | S.E. |  |
| CYA | > 0.54 | - | - | - | -0.237 | 0.078 |  |
| FLU | > 1.7 | - | - | - | -0.898 | 0.409 |  |
| SUL | 0.025 | 0.002 | 0.022 | 0.028 | -4.528 | 0.720 |  |
| THX | 0.0016 | 0.0002 | 0.0011 | 0.0021 | -1.218 | 0.174 |  |
| 48 hour |  |  |  |  |  |  |  |
|  | LD50 |  |  |  | Slope |  |  |
|  | EST | S.E. | LOW | UPP | Honey bee EST<sup>a</sup> | EST | S.E. |
| CYA | > 0.54 | - | - | - | > 0.1055<sup>b</sup> | - | - |
| FLU | > 1.7 | - | - | - | 1.2<sup>c</sup> | -0.751 | 0.374 |
| SUL | 0.0194 | 0.0013 | 0.0169 | 0.022 | 0.146<sup>d</sup> | -3.868 | 0.573 |
| THX | 0.0012 | 0.0002 | 0.0008 | 0.0015 | 0.005<sup>e</sup> | -1.397 | 0.203 |
| 72 hour |  |  |  |  |  |  |  |
|  | LD50 |  |  |  | Slope |  |  |
|  | EST | S.E. | LOW | UPP | EST | S.E. |  |
| CYA | > 0.54 | - | - | - | -0.223 | 0.116 |  |
| FLU | > 1.7 | - | - | - | -0.644 | 0.375 |  |
| SUL | 0.018 | 0.001 | 0.015 | 0.02 | -3.788 | 0.570 |  |
| THX | 0.0012 | 0.0002 | 0.0008 | 0.0015 | -1.397 | 0.203 |  |

<sup>a</sup> Honeybee acute oral LD50 at 48 hours
<sup>b</sup> PRD2013-09 (Health Canada 2013)
<sup>c</sup> PRD2014-20 (Health Canada 2014)
<sup>d</sup> PRD2015-08 (Health Canada 2015)
<sup>e</sup> US EPA (United States Environmental Protection Agency 2017), EFSA (European Food Safety Authority 2015)
Abbrev = A.I. = Active Ingredient, CYA = Cyantraniliprole, FLU = Flupyradifurone, SUL = Sulfoxaflor, THX = Thiamethoxam, EST = Estimate, S.E. = Standard Error, LOW and UPP = Lower and Upper bounds of 95% confidence interval respectively

A dose-response model could not be fitted for the cyantraniliprole data at 48 hours as the highest concentration tested (0.54 μg/bee) produced an average of 41% mortality, thus a 48-hr LD50 could not be calculated. Oral LD50 values of >0.28 μg/bee for bumblebees (*Bombus terrestris*) and >0.11 μg/bee for the European honey bees (*Apis mellifera*) have previously been reported for cyantraniliprole (Dinter and Samel 2014). Health Canada reported the oral LD50 for *A. mellifera* to be >0.1055 μg/bee (Health Canada 2013). While the results from this study, results from Dinter and Samel (2014) and data reported by Health Canada (2013) do not allow us to compare sensitivity across these three bees species, these studies support the view that cyantraniliprole is considerably less toxic to *B. impatiens, B. terrestris*, and *A. mellifera* compared to thiamethoxam and sulfoxaflor.

Although the chronic effects of cyantraniliprole exposure have not been investigated, some inferences can be made from the studies using the closely related compound chlorantraniliprole (Selby et al. 2013). Toxicity tests with chlorantraniliprole produced an estimated oral LD50 of >114 μg/bee at 48 hours for *A. mellifera* (Bassi et al. 2009) and an oral LC50 of 0.013 mg/mL (0.011-0.016 mg/mL 95% C.I.) at 72 hours for *B. terrestris* when exposed via drinking sugar water, however the maximum mortality never reached 100% (79 ± 8% at 40 mg/L) (Smagghe et al. 2013). Sublethal effects of chlorantraniliprole exposure have also been reported for bumblebees. Seven of eight *B. terrestris* microcolonies exhibited sluggish, fatigued behaviour after four weeks of exposure to 0.4 mg/L chlorantraniliprole (100 times less than the maximum field realistic concentration) in pollen, but this effect was not seen for exposure by contact or via drinking sugar water at 0.4 mg/L (Smagghe et al. 2013). However, *B. impatiens* microcolonies fed chlorantraniliprole-treated (0.000615 mg/g Altacor 35 WG) pollen showed no differences in average worker lifespan, amount of pollen consumed, number of days to first oviposition or number of ejected larvae compared to untreated controls (Gradish et al. 2010). In a field test where *B. impatiens* were exposed to chlorantraniliprole sprayed onto weedy turf grass and white clover, colonies showed no impairment in weight gain or the number of queens produced relative to controls, and workers from these colonies did not appear to avoid foraging on treated plots (Larson et al. 2013).

Thiamethoxam was the most toxic of the four active ingredients for *B. impatiens* with an LD50 of 0.0012 μg/bee (Table 2). The LD50 value for thiamethoxam differs substantially from values reported from group feeding trials (of 5-9 bumblebees per cage) using doses ranging from 6.68 pM to 8.55 mM in a 2:1 honey-water ratio (Baines et al. 2017). Interestingly, these authors found high *B. impatiens* worker mortality at both the lowest (58% between 0.039-0.78 pg/μL) and highest treatment concentrations (100% at 1.1 pg/μL), with reduced mortality rates (33-51%) for intermediate concentrations (15.6-312.5 pg/μL). Thus, they classified the dose response curve for thiamethoxam as biphasic, leading to two different LD50 estimates at the low and high peak, 1.88 x 10^-6^ and 8.18 x 10^-6^ μg/bee, respectively (reported values: 22.6 pg/120μL and 98.2 pg/120μL: Baines et al. 2017). As there were substantial differences in the methods used by these authors compared to our study, such as non-standardized dose consumption by individual worker bees (range 55-120 μL consumed), a longer time period for consumption (24 hours) and feeding as a group, further investigation into the effect of methods on the production and comparability of LD50 values should be considered a high priority for risk assessments.

The relatively high toxicity of thiamethoxam in this study is not surprising given the potency of the neonicotinoid class on honey bees and other bumblebee species (Arena and Sgolastra 2014). Recent work suggests dietary exposure to similarly low levels of thiamethoxam (2.4 x 10^-6^ - 1.0 x 10^-5^ mg/mL of sucrose solution) can impact egg development and feeding rates in queens (Baron et al. 2017b), colony initiation success by queens (Baron et al. 2017a), worker learning and memory (Stanley et al. 2015b), worker foraging performance (Stanley et al. 2015a; Stanley and Raine 2016) and male size (Stanley and Raine 2017) in bumblebees. Relatively low neonicotinoid residues levels (≤1.0 x 10^-5^ mg/mL) in the nectar or pollen of treated crops or non-crop flowers have the potential to impact not only individuals, but populations within relatively short periods of exposure. This may be concerning as these, or higher, concentrations have been documented in the nectar and pollen of crop flowers or wild plants in Canada (Tsvetkov et al. 2017), Europe (David et al. 2016; Woodcock et al. 2017) and the United States (Long and Krupke 2016; Stewart et al. 2014).

Sulfoxaflor was less toxic than thiamethoxam to *B. impatiens* but had substantially lower LD50 values than cyantraniliprole and flupyradifurone (Table 2). Variation in binding site affinity may be one of the reasons why sulfoxaflor has a different toxicity than thiamethoxam. While sulfoxaflor can bind to the low and high affinity sites on nAChR, it does not bind as strongly to the low affinity site compared to neonicotinoid insecticides (Sparks et al. 2013). Bumblebee (*B. terrestris*) colonies raised from wild caught queens and exposed to 0.005 mg/L of sulfoxaflor for two weeks in the laboratory before being placed out in parkland produced no queens and significantly fewer males compared to untreated control colonies (Siviter et al. 2018). Sulfoxaflor exposure can also affect the number of eggs laid by *B. terrestris* workers in queenless microcolonies, leading to a lower likelihood of producing larvae (Siviter et al. 2020).

In contrast, laboratory tests revealed no evidence of sulfoxaflor exposure impairing *B. terrestris* worker performance in either olfactory learning (using proboscis extension reflex (PER) conditioning) or working memory (using a radial-arm maze) assays (Siviter et al. 2019). These results suggest that impacts on bumblebees can be detected even after a short exposure period at low concentrations. To our knowledge, no studies have yet assessed impacts for bumblebees of chronic exposure to sulfoxaflor over periods longer than 2 weeks. This would be a useful area of future research, to fully characterize effects to increase risk assessment accuracy.

### 3.4 Sublethal responses

Significant evidence of abnormal behavioural responses was found in trials where workers were exposed to flupyradifurone. Exposure to the highest concentration of flupyradifurone (0.17 mg/mL) resulted in 44% mortality at 48 hours, and sublethal effects indicative of neurotoxic impacts (Figure 1). Four hours after pesticide ingestion we found a strong positive correlation between flupyradifurone dose and the proportion of bees displaying abnormal behaviours (r = 0.8, *p* < 0.001: Figure 1A). This high incidence of sublethal effects four hours after pesticide ingestion, followed by the subsequent decrease in proportion of bees exhibiting abnormal behaviour at subsequent time points (Figs 1B-D), is likely explained by the fact that flupyradifurone reversibly binds to the nAChR (Jeschke et al. 2015) allowing some affected bees to recover. However, it could also be that those individual bees most strongly affected by exposure died ahead of subsequent mortality checks, which is supported by the increase in mortality over time for all concentrations. A relatively strong positive correlation between flupyradifurone dose and the proportion of bees displaying abnormal behaviours was still evident 24 hours after pesticide ingestion (r = 0.55, p < 0.001: Fig 1B), but at 48 hours the relationship weakened (r = 0.2, p = 0.25: Fig 1C), and by 72 hours all surviving bees showed no behavioural abnormalities (Fig 1D). Bumblebees exposed to the highest dose showed higher mortality than the lowest dose (44% vs. 27%) for all trials at 72 hours. The lower survivorship of bees exposed to the highest dose suggests they were less likely to recover from sublethal effects than bees exposed to lower concentrations. Indeed, 92% of bees (35/38) exposed to the highest concentration expressed abnormal behaviour, and 54% of those bees (19/35) showing sublethal effects ultimately survived 72 hours. At the lowest exposure level, 41% of bees (16/39) expressed abnormal behaviour and 68% of those bees (11/16) survived 72 hours.

**Figure 1.**
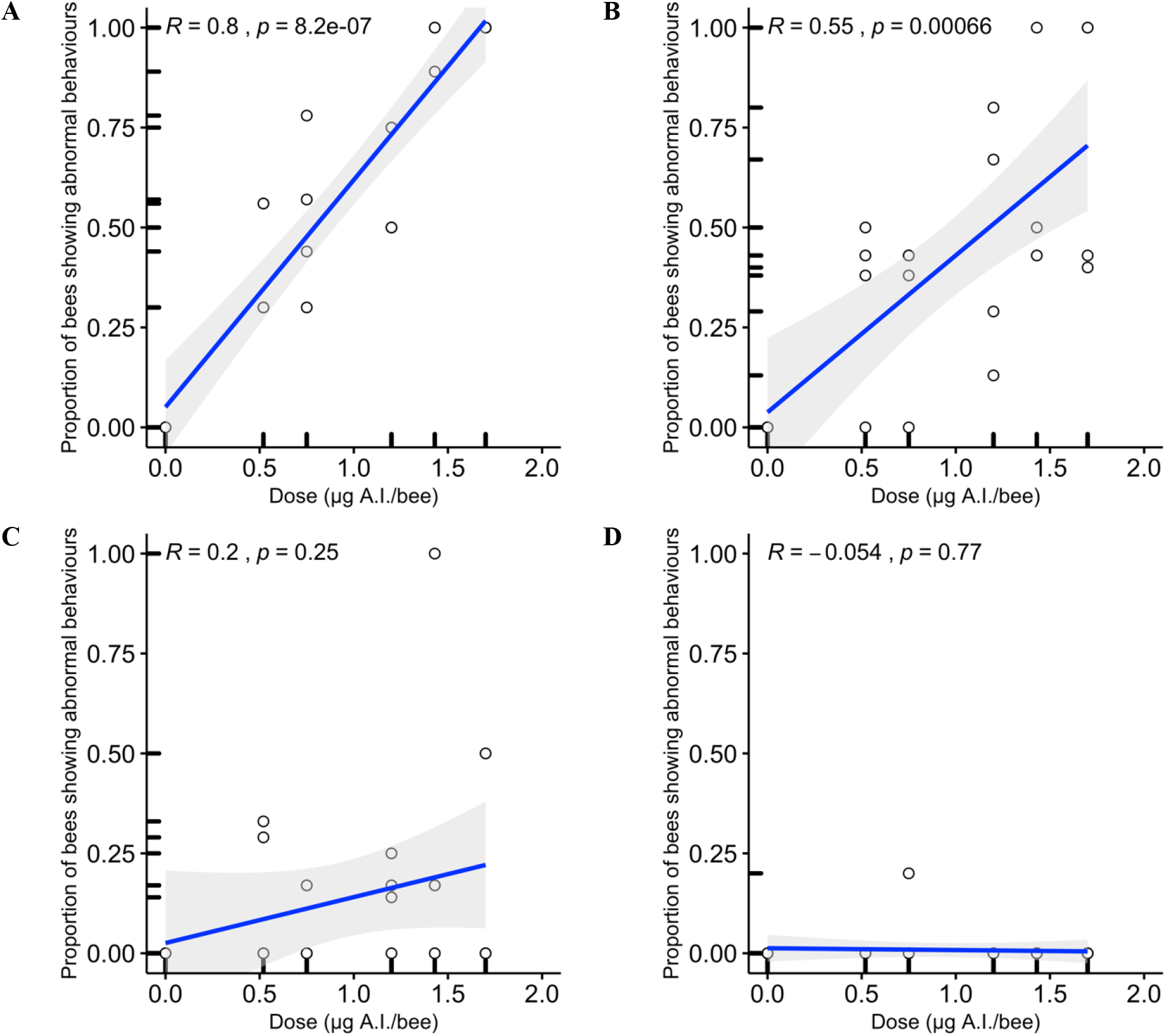
The proportion of bees displaying abnormal behaviours (sublethal effects) from flupyradifurone exposure across the range of tested doses at 4 (A), 24 (B), 48 (C) and 72 (D) hours post consumption. Kendall rank correlation lines of best fit are displayed as blue lines with grey shading indicating 95% confidence intervals. Rank coefficient and *p* value are given in top left corner.

Currently few studies have been conducted on the potential impacts of flupyradifurone on bees, which is surprising given that it is registered for a variety of flowering and non-flowering crops using foliar, soil drench and seed coating applications (European Commission 2016; Health Canada 2014). Sublethal effects of flupyradifurone exposure for honey bees *(Apis* spp.) include impairments of motor abilities (Hesselbach and Scheiner 2019), gustatory responses to sucrose (Hesselbach and Scheiner 2018) and learning and memory performance (Hesselbach and Scheiner 2018; Tan et al. 2017). Acute oral exposure to flupyradifurone (1.2 □ μg/bee) was sufficient to impair *A. mellifera* worker motor abilities, similar to behavioural changes resulting from much lower level exposure (4 ng/bee) of the neonicotinoid, imidacloprid (Hesselbach and Scheiner 2019). Acute oral exposure to 1.2□μg of flupyradifurone was also sufficient to adversely affect the responsiveness of individual *A. mellifera* pollen or nectar foragers to sucrose across a range of concentrations, and also to inhibit their subsequent performance in olfactory learning (PER) assays (Hesselbach and Scheiner 2018). Flupyradifurone exposure can also affect olfactory learning performance in Asian honey bees (*Apis cerana*) whether consumed as either larvae or adults (Tan et al. 2017). Larvae fed the lower dose (0.033 μg/day for 6 days) did not differ in survival from unexposed controls, but those fed a higher dose (0.33 μg/day for 6 days) were less likely to survive to the sealed cell phase and emerge as adults compared to controls. Bees exposed to the low dose as larvae exhibited slower olfactory learning in a PER conditioning assay than controls as adults (Tan et al. 2017). Workers exposed to flupyradifurone as adults had survival rates of 80% when fed 0.066 μg/day and 50% when fed 0.66 μg/day (high dose) for three days (Tan et al. 2017). Olfactory learning performance was adversely affected for surviving bees exposed to either the low or high dose of flupyradifurone (two, rather than one trial learning in the PER assay) compared to control bees (Tan et al. 2017). These results suggest that flupyradifurone has a stronger effect on learning if worker bees are exposed as larvae rather than as freshly emerged adults. The lowest dose tested in this study was 0.0165 μg/bee, compared to a high dose of 0.165 μg/bee (Tan et al. 2017) and 12 ng – 1.2 μg/bee (Hesselbach and Scheiner 2018; 2019). Adding our results to those reported by others suggests a range of sublethal effects, from obvious central nervous system malfunction to more subtle learning and memory impairment. However, as many of these effects occurred at levels substantially above exposure levels expected under field conditions, more work is needed to determine the potential impacts on bumblebees exposed to lower levels of flupyradifurone.

### 3.5 Comparing toxicity between Apis mellifera and Bombus impatiens

A comparison of the oral LD50 values for *B. impatiens* from the present study against published LD50 values for *A. mellifera* (Table 2) highlights that honey bees are not always the most sensitive species to insecticide exposure. Based on the 48-hr LD50s reported for honey bees, sulfoxaflor and thiamethoxam are more acutely toxic to bumblebee workers (*B. impatiens*; Table 2): sulfoxaflor and thiamethoxam are approximately 7.5 and 4.2 times more toxic to bumblebees than to honey bees respectively. Given the uncertainties that exist around the relative toxicity of compounds to different bee species compared to the honey bee model for ecotoxicological testing, some regulators (e.g. European Food Safety Authority) include a suggested 10x safety factor − i.e. dividing the LD50 values for honey bees by 10 on the assumption that other bee species might be 10x more sensitive to these pesticides (Arena and Sgolastra 2014; Heard et al. 2017). This 10x safety factor would be sufficiently protective for all four insecticides tested on *B. impatiens* in the present study (Table 2).

Flupyradifurone is estimated to be 1.4 times more toxic to honey bees than bumblebees, but this difference could be greater as the bumblebee LD50 was not established at the highest level of exposure (1.2 μg/bee) in the present study. It is not clear whether there are differences in sensitivity between these species for cyantraniliprole, because the actual LD50 values could not be determined. The LD50 for cyantraniliprole is reported as >0.54 μg/bee in the present study and the reported honey bee value is >0.1055 μg/bee (Health Canada 2013). These data should be used to inform reviews of current risk assessment practices for pesticide exposure for insect pollinators to ensure any safety factors used are sufficiently protective to cover the full range of differential sensitivities among insect pollinator taxa.

### 3.6 Risk Assessment

The EPPO (European and Mediterranean Plant Protection Organization 2010) hazard quotient (HQ) classified all active ingredients as a moderate or high hazard (HQ ≥50: Table 3). This hazard quotient method allows an initial estimation of how much risk a honey bee may face from the application of an insecticide. Regardless of application rate, flupyradifurone (mean: 744 mL/ha, range: 614-875 mL/ha) was always classified as moderate-risk, whereas thiamethoxam (mean: 690 mL/ha, range: 629-750 mL/ha application rates) was always high-risk. Sulfoxaflor (mean: 275 mL/ha, range: 224-326 mL/ha application rates) was predominately high-risk for average and minimum application rates but increased to entirely high-risk for maximum application rates (Table 3). Cyantraniliprole (mean: 773 mL/ha, range: 606-940 mL/ha application rates) showed the greatest variation in risk class based on application rate. Moving from minimum to maximum application rates produced a shift from 100% moderate to 21% high-risk. Thus, for some active ingredients knowledge about the likelihood that particular application rates will be required or used will inform estimates about which risk classification is most relevant. As these application rates all follow manufacturer recommendations, these represent real world exposure scenarios. Higher application rates may be used by growers when either more robust treatments are triggered or if farmers employ a more conservative spray regime (i.e. spray at pest presence rather than when particular economic thresholds are reached).

**Table 3.** Risk classification of active ingredients using the hazard quotient (HQ) risk equation. Data show the percentage of times an insecticide was classified as either a moderate or high HQ using three different application rates. Risk levels for the HQ are low (<50), moderate (50-2500) or high (>2500) (European and Mediterranean Plant Protection Organization 2010).

| A.I. | Application Rate (mL/ha) |  |  |  |  |  |
| --- | --- | --- | --- | --- | --- | --- |
|  | Minimum |  | Average |  | Maximum |  |
|  | Moderate HQ | High HQ | Moderate HQ | High HQ | Moderate HQ | High HQ |
| **CYA**<sup>a</sup> | 100% | 0% | 99% | 1% | 79% | 21% |
| **FLU**<sup>b</sup> | 100% | 0% | 100% | 0% | 100% | 0% |
| **SUL**<sup>c</sup> | 6% | 94% | 6% | 94% | 0% | 100% |
| **THX**<sup>d</sup> | 0% | 100% | 0% | 100% | 0% | 100% |

<sup>a</sup> Application rates for Minimum value follow a minimum application amount for all pests that the product could be used against for Dupont Benevia Insecticide, Dupont Exirel Insecticide, Dupont Okina Insect Control, Dupont Verimark Insecticide, Dupont™ HGW86 200 SC Potato Seed Treatment Insecticide. Averages were calculated using the median application rates for all pests against which the product could be used. Maximum was calculated using the highest application amount for all pests that the product could be used against. 773, 606, and 940 mL/ha are the mean average, minimum and maximum application rates, respectively (see Table S3 for all mL/ha values).
<sup>b</sup> Application rates for Minimum value follow a minimum application amount for all pests that the product could be used against for Sivanto Prime Insecticide. Averages were calculated using the median application amount for all pests that the product could be used against. Maximum was calculated using the highest application amount for all pests that the product could be used against. 744, 614, and 875 are the mean average, minimum and maximum application amounts, respectively (See Table S3 for all mL/ha values).
<sup>c</sup> Application rates for Minimum value follow a minimum application amount for all pests that the product could be used against for Twinguard Insecticide and Closer Insecticide. Averages were calculated using the median application amount for all pests that the product could be used against. Maximum was calculated using the highest application amount for all pests that the product could be used against. 275, 224, and 326 are the mean average, minimum and maximum application amounts, respectively (See Table S3 for all mL/ha values).
<sup>d</sup> Application rates for Minimum value follow a minimum application amount for all pests that the product could be used against for Minecto Duo 40WG. Averages were calculated using the median application amount for all pests that the product could be used against. Maximum was calculated using the highest application amount for all pests that the product could be used against. 690, 629, and 750 are the mean average, minimum and maximum application amounts, respectively (See Table S3 for all mL/ha values).
Abbrv = A.I. = Active Ingredient, CYA = Cyantraniliprole, FLU = Flupyradifurone, SUL = Sulfoxaflor, THX = Thiamethoxam, HQ = Hazard Quotient

The standard risk approach (SRA) and fixed dose risk approach (FDRA) equations categorized all active ingredients into low, moderate, high or very high-risk categories (Sanchez-Bayo and Goka 2014) (Table 4). On average, pollen residues were 668, 1479, 12932, and 17 ng/g and mean nectar residues were 78, 178, 551, and 2 ng/g for cyantraniliprole, flupyradifurone, sulfoxaflor, and thiamethoxam, respectively (Tables S3, S4). Using the FDRA equation, the majority of cyantraniliprole calculations were categorised as low-risk for pollen (92%) and nectar (100%) and fewer as moderate-risk (pollen, 8%, Table 4). Comparatively, the cyantraniliprole SRA calculations placed a higher percentage within the moderate and high-risk categories (pollen 8% and nectar 20% in each category, Table 4). Flupyradifurone was categorized as high-risk most often for nectar (85%) using the SRA, but pollen was predominately low-risk (57%), whereas the FDRA most commonly placed pollen (57%) and nectar (62%) in the low-risk category (Table 4). Flupyradifurone residues in pollen also appeared in the moderate (29%) and high (14%) risk FDRA categories, and the rest of the nectar calculations were categorized as moderate (38%) risk. Sulfoxaflor was predominately within the high-risk SRA categories for pollen (70%) and nectar (100%), and high-risk FDRA categories for pollen (60%) and nectar (75%) (Table 4). Thiamethoxam was always placed in the high-risk category for pollen and nectar using the SRA and for nectar using the FDRA (Table 4). Pollen was predominately categorized as low (67%) or moderate (33%) risk in FDRA calculations (Table 4).

**Table 4.** Risk classification of active ingredients using the standard risk approach (SRA) and the fixed dose risk approach (FDRA) equations. Data show the percentage of times an insecticide was low, moderate or high scoring under each risk assessment scenario. Pollen and nectar risks were examined separately using residue data specific to each matrix. Risk levels for SRA: Low <1%, Moderate = 1-5%, High >5% (Sanchez-Bayo and Goka 2014). Risk levels for FDRA: Low 7 -≥ 60 days, moderate between 2-7 days High ≤2 days (Sanchez-Bayo and Goka 2014).

| A.I. | Standard Risk Approach |  |  |  |  |  | Fixed Dose Risk Approach |  |  |  |  |  |
| --- | --- | --- | --- | --- | --- | --- | --- | --- | --- | --- | --- | --- |
|  | Low Risk |  | Moderate Risk |  | High Risk |  | Low Risk |  | Moderate Risk |  | High Risk |  |
|  | Pollen | Nectar | Pollen | Nectar | Pollen | Nectar | Pollen | Nectar | Pollen | Nectar | Pollen | Nectar |
| CYA | 85% | 60% | 8% | 20% | 8% | 20% | 92% | 100% | 8% | 0% | 0% | 0% |
| FLU | 57% | 15% | 14% | 0% | 29% | 85% | 57% | 62% | 29% | 38% | 14% | 0% |
| SUL | 20% | 0% | 10% | 0% | 70% | 100% | 20% | 0% | 20% | 25% | 60% | 75% |
| THX | 0% | 0% | 0% | 0% | 100% | 100% | 67% | 0% | 33% | 0% | 0% | 100% |

<sup>a</sup> Mean pollen residues were 668, 1479, 12932 and 17 ng/g for CYA, FLU, SUL and THX respectively. See Table S4 for specific residue information.
<sup>b</sup> Mean nectar residues were 78, 178, 551 and 2 ng/g for CYA, FLU, SUL and THX respectively. See Table S4 for specific residue information.
Abbrev = A.I. = Active Ingredient, CYA = Cyantraniliprole, FLU = Flupyradifurone, SUL = Sulfoxaflor, THX = Thiamethoxam.

While the risk categories for each active ingredient varied little, if at all, across application rates when using the HQ approach (Table 3), risk categorization patterns were considerably more variable when using either the SRA or FDRA (Sanchez-Bayo and Goka 2014) (Table 4). This variation in risk categorization using the SRA or FDRA may be helpful for regulators to consider so they can more effectively tailor future pollinator risk assessments to real variation in the exposure to active ingredients found in pollen and nectar across a variety of crops as this information is reported for registration purposes (Health Canada 2013; 2014; 2015). Determining which crops, and even which application periods, pose the lowest risks to bees will help as pest management practices are refined to minimize impacts of pesticide exposure when and where these agrochemicals are used.

## 4 CONCLUSION

This study provides acute oral toxicity data for the Common Eastern bumblebee, *Bombus impatiens* when individually exposed to four different systemic insecticides across a range of concentrations: cyantraniliprole, flupyradifurone, sulfoxaflor and thiamethoxam. As each insecticide is registered for use in Canada, the United States and the European Union, risks of exposure for bumblebees, and other insect pollinators, could be widespread. However, because only *B. impatiens* was tested, which is native to eastern Canada and the eastern United States, caution should be taken when extrapolating from this study to other bumblebee species and other bee taxa. Based upon our assessment, thiamethoxam (the neonicotinoid class representative) appears to be the most acutely toxic to *B. impatiens*, followed by sulfoxaflor (sulfoximine), flupyradifurone (butenolide) and cyantraniliprole (diamide). More research should be conducted on the precise modes of action and metabolic processes that break down these insecticides to provide a better understanding of their differential toxicity. Our results also suggest the need for additional work to understand the impacts of chronic exposure to sublethal concentrations, as earlier research has shown potential for impacts on bee populations following exposure to thiamethoxam or other neonicotinoids (Baron et al. 2017a; Rundlöf et al. 2015; Woodcock et al. 2017). Greater understanding of why these four systemic insecticides have such different impacts on this common bumblebee species could lead development towards biochemical scaffolds that are less harmful to major beneficial insect groups while effectively controlling targeted insect pests. Based upon this research, it might be advisable to investigate other butenolide and diamide active ingredients for use where systemic insecticides might be the best pest control option to minimise potential impacts for bees.

Current guidelines suggesting a 10x safety factor (Arena and Sgolastra 2014; Heard et al. 2017) capture the variation in toxicity for these four active ingredients. The present study showed that *B. impatiens* was more sensitive than *A. mellifera* to thiamethoxam and sulfoxaflor. Although the majority of bumblebee insecticide toxicity studies to date have focused on *B. terrestris* (a European species) and have suggested that *A. mellifera* was more sensitive than *B. terrestris*, the present study indicates that this may not be the relationship for all bumblebee species (Arena and Sgolastra 2014). It is important to consider which beneficial insects will be performing crop pollination services or included as a part of integrated pest management (IPM) strategies, and choose pesticides that simultaneously minimize harm to the beneficial insects and maximize effective control of insect pests. Ensuring that toxicity tests are done on native bumblebee species would help build more locally applicable toxicity rankings which should be incorporated into appropriate risk assessments. Further research should be also conducted regarding the impacts of all exposure routes for solitary bees (including exposure from soil for ground-nesting species (Chan et al. 2019)) to determine if risks change based upon differences in ecology and life history.

At least some application scenarios for each active ingredient were categorised as moderate or high-risk regardless of the risk assessment equation applied. While the EPPO (European and Mediterranean Plant Protection Organization 2010) HQ methods provides basic information on potential risk levels, incorporating the likelihood that a pollinator comes into contact with pesticides residues using the SRA or FDRA (Sanchez-Bayo and Goka 2014) approaches revealed substantial variability in risk categories for each active ingredient and that hazard levels can be altered based upon the food source consumed by bumblebees. Moving forward, risk assessments should use more nuanced calculations that will increase our understanding of risk for not only bumblebees, but also for other wild pollinators that may also be exposed. Risk assessment of insecticide exposure for wild bee populations is a burgeoning field of research and will continue to grow as frameworks of management practices for bees develop and the importance of wild pollinators for sustainable agricultural production becomes even clearer.

## Notes

#### Summary of Updates

Error in Table 2 has identified and corrected

